## Supplementary Data for "Inhibitory circuits control leg movements during *Drosophila* grooming"

### Supplementary Figures and Videos

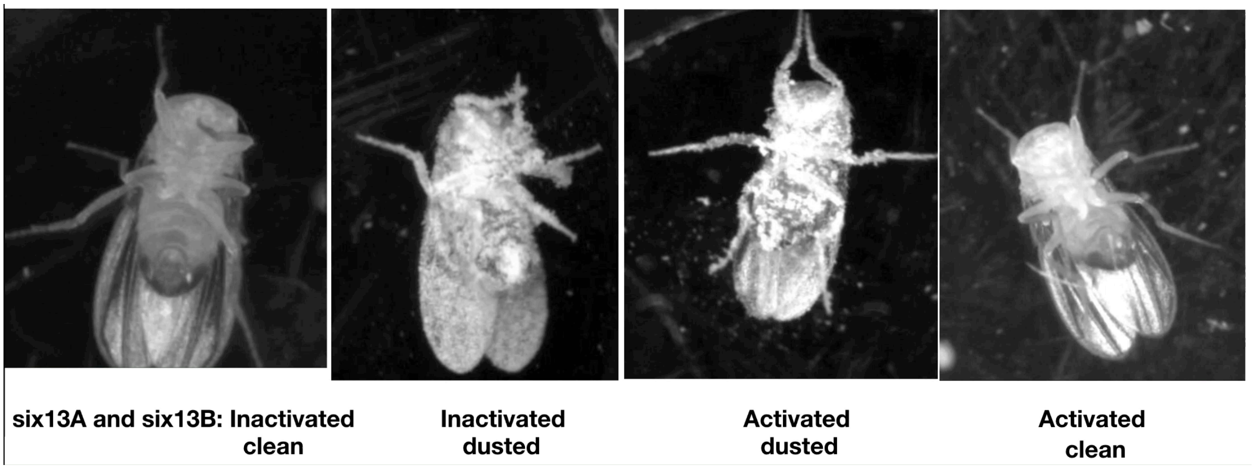

**<u>Figure 1—Video 1</u>**. Manipulating activity of six 13A neurons and six 13A neurons in headless flies. Inactivation: Legs locked in flexion in clean and dust covered flies. Activation: legs extended in clean and dust covered flies.

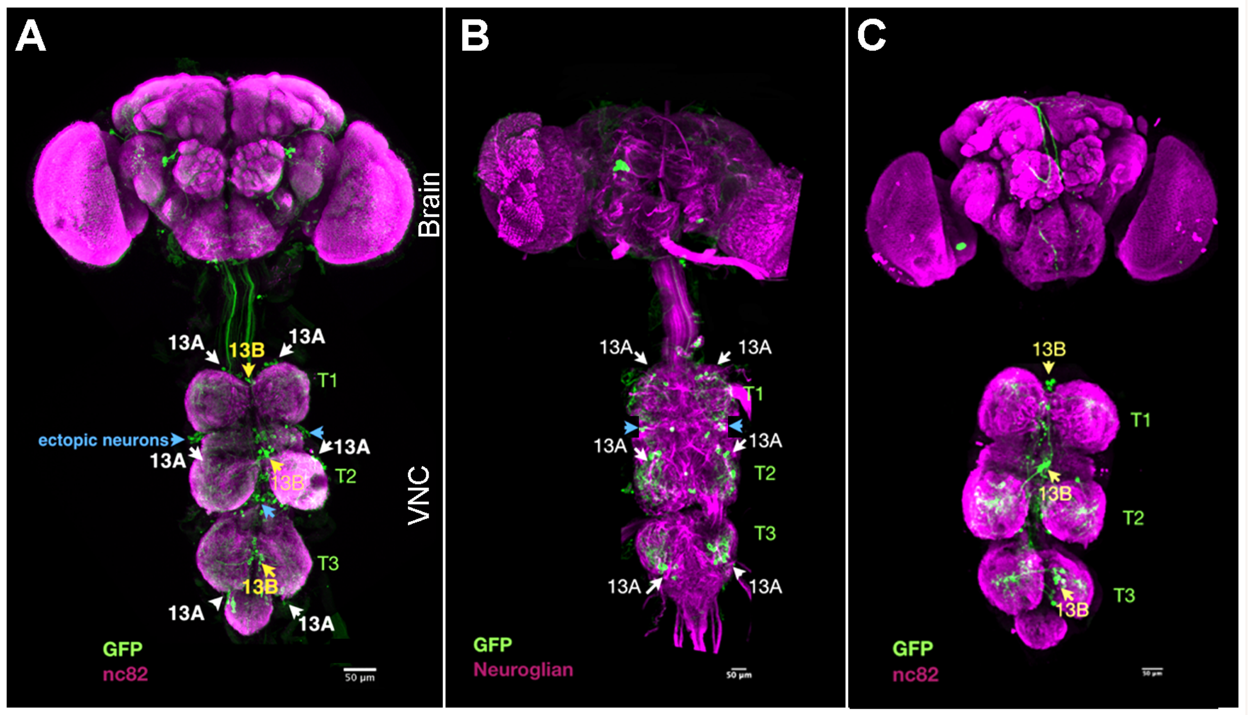

**Figure 1—figure supplement 1. Expression pattern in the central nervous system of various lines used for behavior experiments**

(A) R35G04-DBD and GAD1-AD Split GAL4 combination labels approximately 6-7 13A neurons and 3 13B neurons per thoracic hemisegment. Ectopic expression is observed in a few neurons in the brain, and in the ventral nerve cord (VNC) in about 9 neurons per hemisegment (possibly 13A neurons with posterior cell bodies [42,85],or 3B neurons) and 4 neurons per hemisegment (possibly 0A) within the Accessory Mesothoracic neuropil (AMNp) and T2 midline.

(B) R35G04-DBD and Dbx-AD Split GAL4 combination specifically labels a subset of 13A neurons already included in the R35G04-DBD and GAD1-AD Split GAL4 line, thereby isolating 13A neurons without labeling 13B neurons. It also labels approximately 3 neurons per hemisegment with posterior cell bodies.

(C) R11B07-DBD and GAD1-AD Split GAL4 combination labels three 13B neurons. It occasionally labels two ectopic ascending neurons per hemisegment.

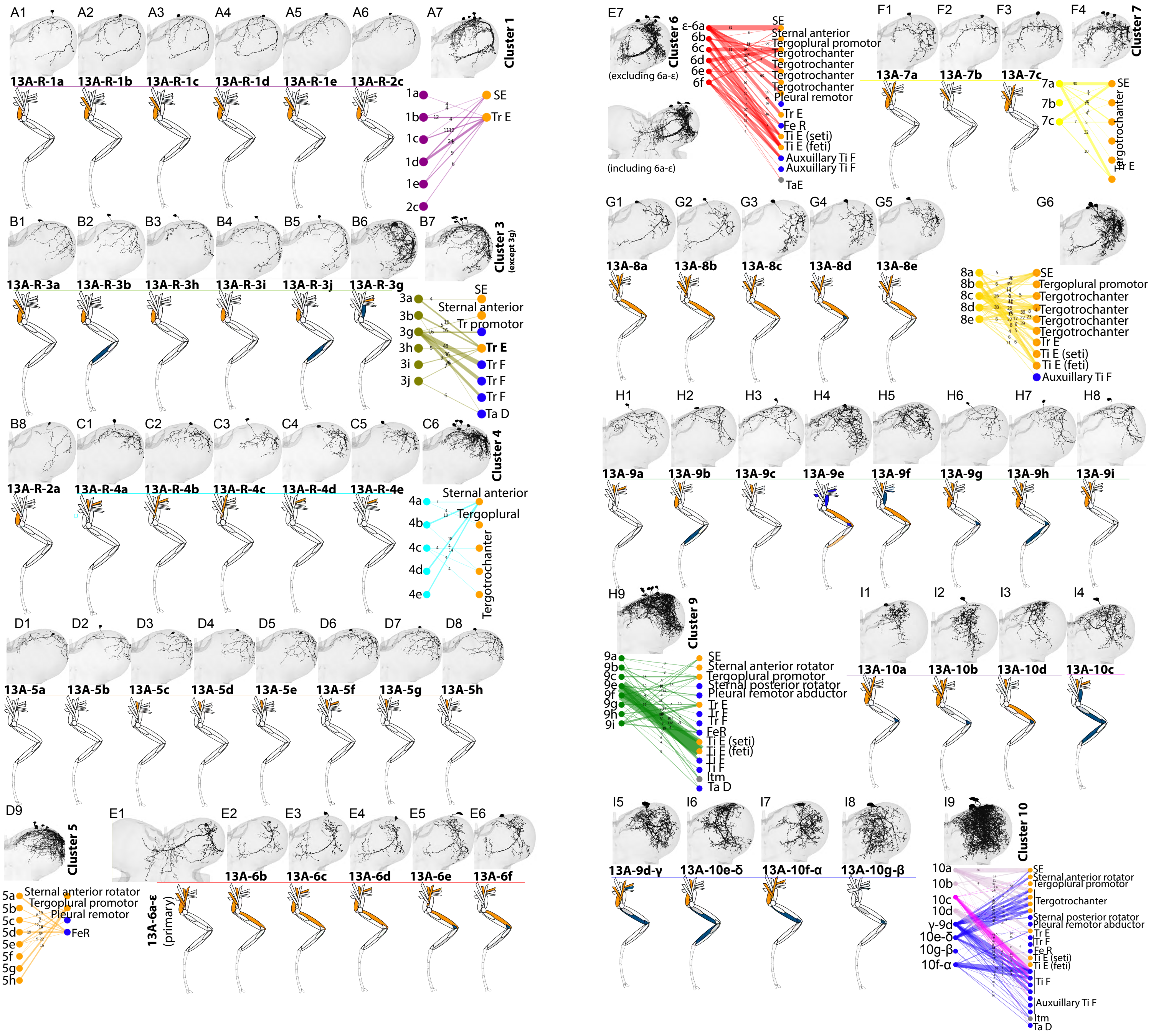

**Figure 2—figure supplement 1. Spatial Map and Connectivity of Premotor 13A Neurons**

(A1-A6) Top: EM reconstruction showing morphology of individual 13A neurons in cluster-1 in the right hemisegment of prothoracic ganglion (T1) region of VNC. Bottom: Leg schematic illustrating muscles innervated by the MNs inhibited by Cluster-1 13A neurons.

(A7) Top: EM reconstruction showing morphology of all cluster-1 13A neurons combined. Bottom: Connectivity matrix showing cluster 1 13A neurons and their MN connections. These neurons connect to Sternotrochanter extensor (SE) and trochanter extensor (Tr E) motor neurons. Edges represent synaptic weight.

(B1-B6) Top: EM reconstruction showing morphology of individual Cluster-3 13A neurons. Bottom: Leg schematic highlighting muscles innervated by the MNs inhibited by Cluster-3 13A neurons.

(B7) Top: Morphology of Cluster-3 13A neurons combined except 13A-R-3g. Bottom: Connectivity matrix showing connections between cluster-3 13A neurons and MNs. Tr =trochanter, Tr E = trochanter extensor, Tr F = trochanter flexor, Ta D= tarsus depressor.

(B8) Top: Morphology of Cluster-2 13A neuron. Bottom: Leg schematic for muscles innervated by the Cluster-2 13A neuron.

(C1-C5) Top: EM reconstruction showing morphology of individual Cluster-4 13A neurons. Bottom: Leg schematic highlighting muscles innervated by the MNs inhibited by Cluster-4 13A neurons.

(C6) Top: Morphology of Cluster-4 13A neurons combined. Bottom: Connectivity matrix showing connections between cluster-4 13A neurons and MNs.

(D1-D8) Top: EM reconstruction showing morphology of individual Cluster-5 13A neurons. Bottom: Leg schematic highlighting muscles innervated by the MNs inhibited by Cluster-5 13A neurons

(D9) Top: Morphology of Cluster-5 13A neurons combined. Bottom: Connectivity matrix showing connections between cluster-5 13A neurons and MNs. Fe R = femur reductor.

(E1-E6) Top: EM reconstruction showing morphology of individual Cluster-6 13A neurons. Bottom: Leg schematic highlighting muscles innervated by the MNs inhibited by Cluster-6 13A neurons.

(E7) Left: Morphology of Cluster-6 13A neurons combined. Right: Connectivity matrix showing connections between cluster-6 13A neurons and MNs.

(F1-F3) Top: EM reconstruction showing morphology of individual Cluster-7 13A neurons. Bottom: Leg schematic highlighting muscles innervated by the MNs inhibited by Cluster-7 13A neurons.

(F4) Top: Morphology of Cluster-7 13A neurons combined. Bottom: Connectivity matrix showing connections between cluster-7 13A neurons and MNs.

(G1-G5) Top: EM reconstruction showing morphology of individual Cluster-8 13A neurons. Bottom: Leg schematic highlighting muscles innervated by the MNs inhibited by Cluster-8 13A neurons.

(G6) Top: Morphology of Cluster-7 13A neurons combined. Bottom: Connectivity matrix showing connections between cluster-7 13A neurons and MNs.

(H1-H8) Top: EM reconstruction showing morphology of individual Cluster-9 13A neurons. Bottom: Leg schematic highlighting muscles innervated by the MNs inhibited by Cluster-9 13A neurons.

(H9) Top: Morphology of Cluster-9 13A neurons combined. Bottom: Connectivity matrix showing connections between cluster-9 13A neurons and MNs.

(I1-I8) Top: EM reconstruction showing morphology of individual Cluster-10 13A neurons. Bottom: Leg schematic highlighting muscles innervated by the MNs inhibited by Cluster-10 13A neurons.

(I9) Top: Morphology of Cluster-10 13A neurons combined. Bottom: Connectivity matrix showing connections between cluster-10 13A neurons and MNs.

Protractors/levators/extensors in orange and retractors, depressors, flexors in blue.

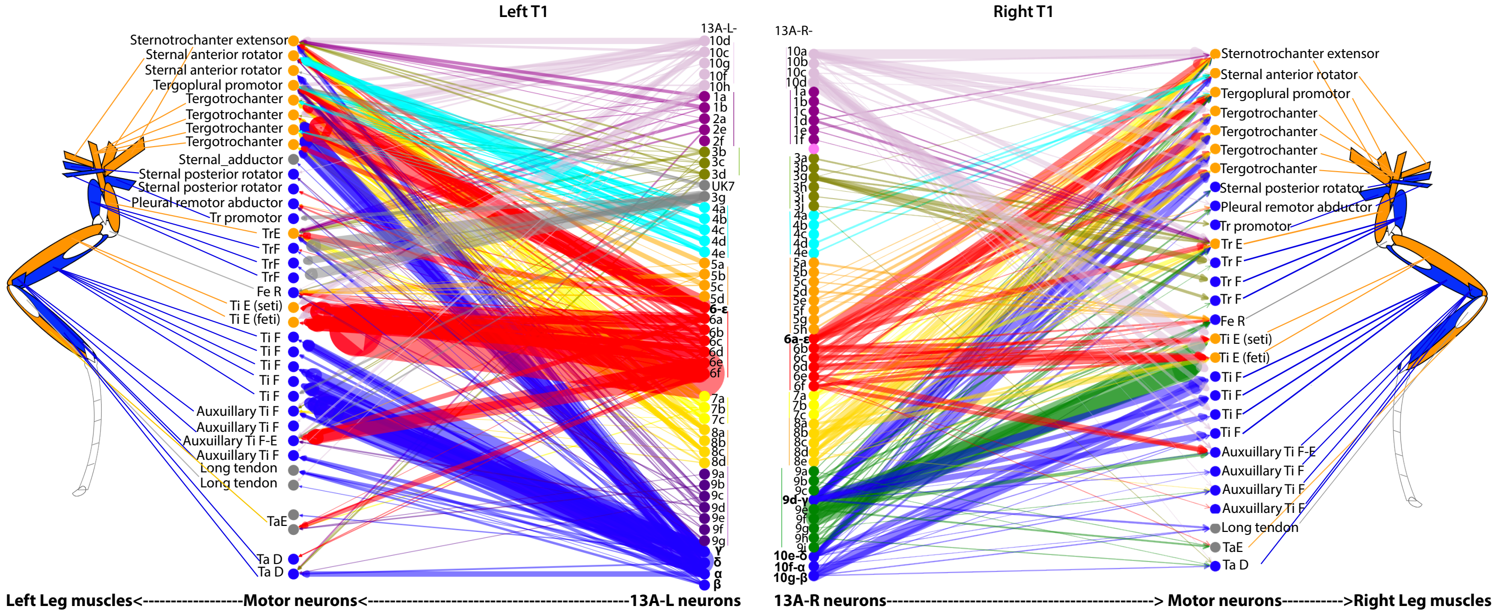

**Figure 2—Figure Supplement 2. Connectivity Matrix of 13A Neurons and the Motor Neurons Showing Left-Right Comparison and Spatial Map.** The 13A neurons that belong to the same anatomical cluster connect to same set of motor neurons (MNs). 13A neurons that have similar morphology are shown in the same color. Clusters that contain similar neurons on the *right and the left* are shown in one color. Clusters are matched as groups, but individual neurons within a cluster are not necessarily one-to-one homologs across sides. The edge width between 13A neurons and motor neurons corresponds to the normalized synaptic weight. Each cluster on the right and the left connects to the same set of MNs. Synaptic weights on the left[15] are greater than on the right, but the connections and preferred partners are the same. The leg schematic shows the muscles that these MNs innervate. Protractors/levators/extensors: orange, retractors, depressors, flexors: blue.

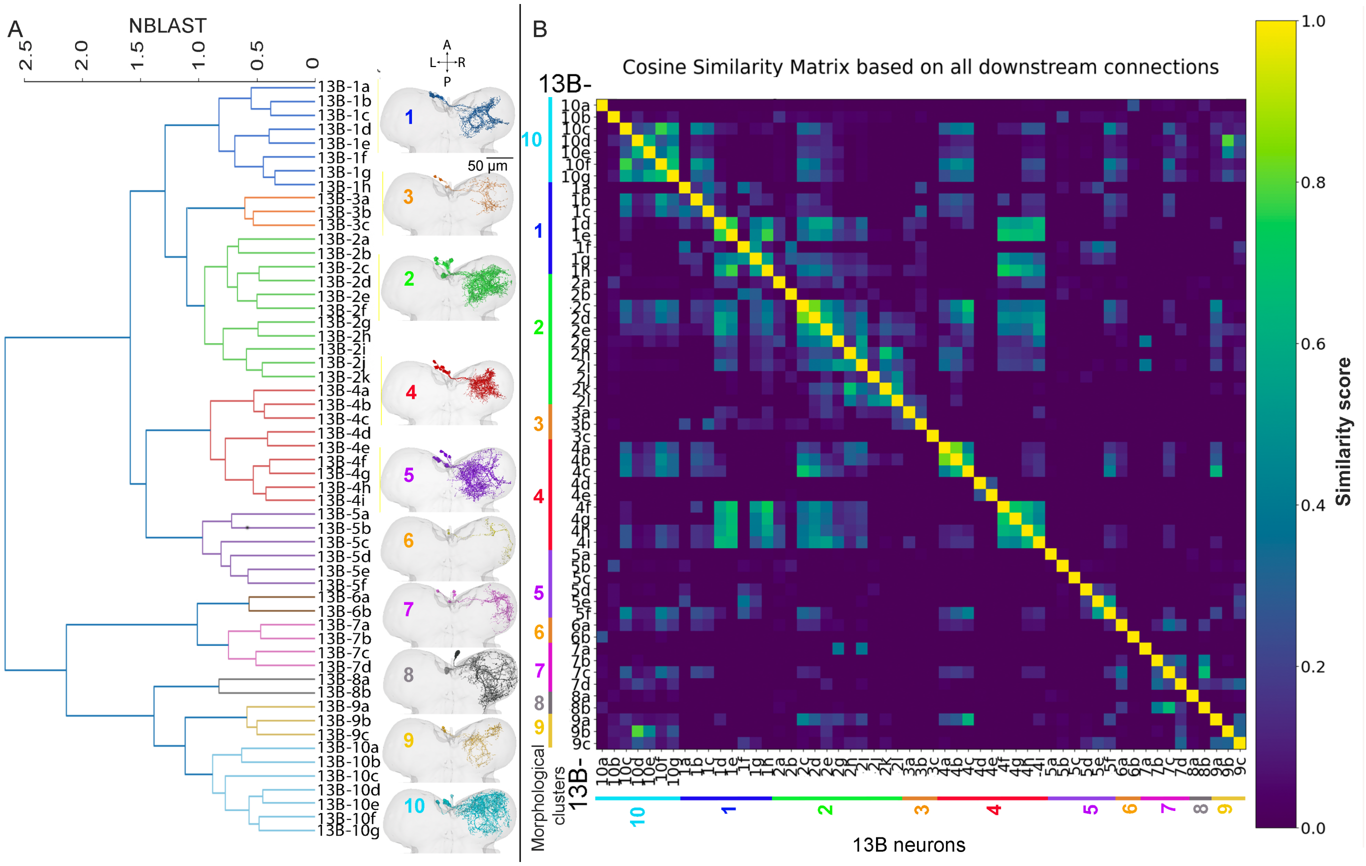

**Figure 2—figure supplement 3. Anatomical classification of 13B neurons.**

(A) Hierarchical clustering of 13B hemilineage in the right (R) T1 based on NBLAST similarity scores. Each cluster is depicted in a different color. The neurons having similar morphology represent one cluster. For example, all neurons in the 13B-1 series have similar morphology and contain 8 neurons 13B-1(a-h). Right panel shows morphologies of neurons in each cluster in T1. Cell bodies are towards left side of the midline but all axonal and dendritic projections are towards right T1.. Each subplot depicts a distinct cluster, with different colors indicating different clusters. A= anterior, P=posterior, L= left, R=right. Ventral side is up.

(B) Cosine similarity graph showing the pairwise similarity between 13B neurons based on their connections to all downstream neurons. 13B neurons are named based on the anatomical clusters obtained with NBLAST as described above.

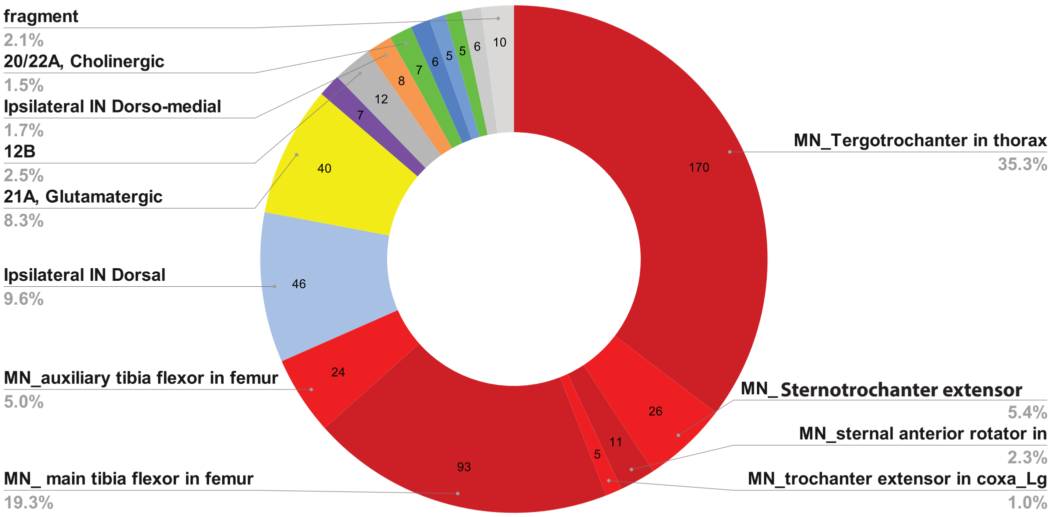

**Figure 2—figure supplement 4. Neurons Downstream of a Primary 13A-10f-α Neuron.** Motor neurons (red) constitute 58.4% of the total downstream synapses. Other downstream connections include glutamatergic hemilineages, 21A, 24B (yellow), cholinergic hemilineages 7B, 3A, 20A (green), GABAergic hemilineages 12B (gray), and other unknown neurons (blue, orange).

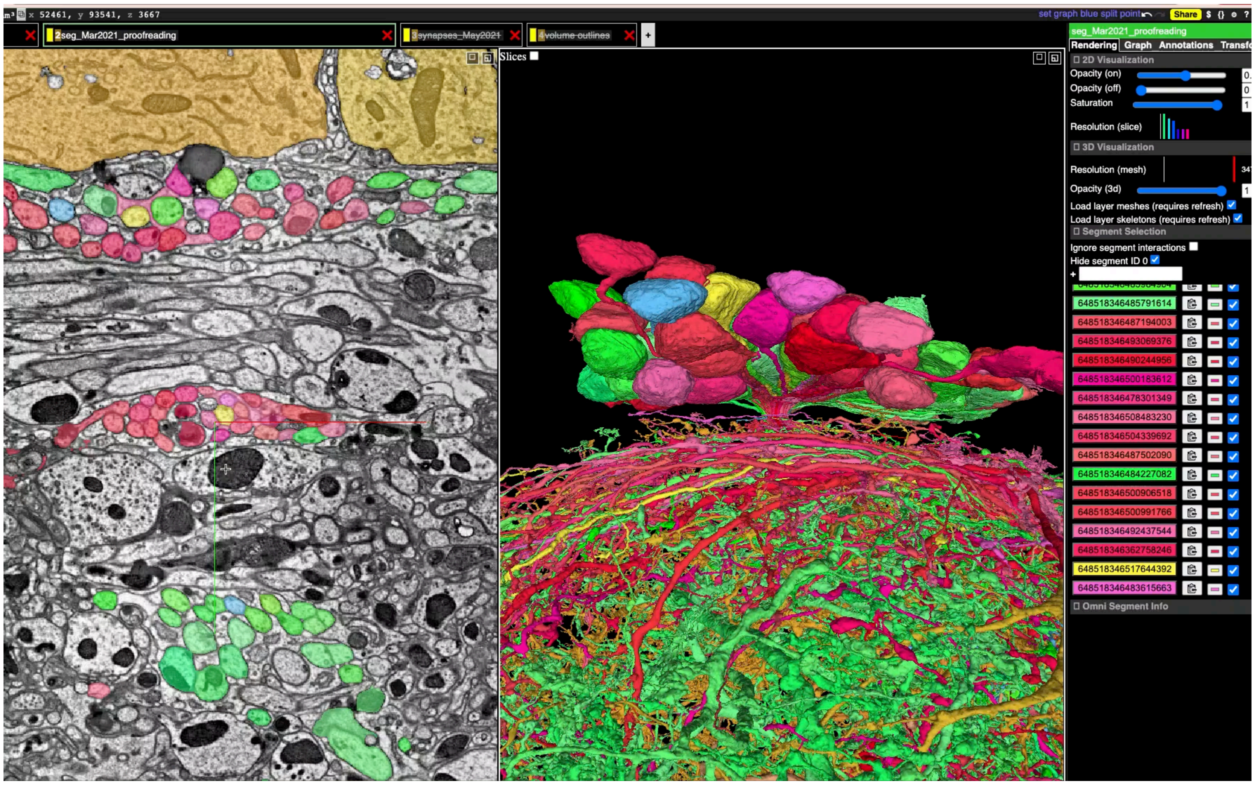

**Figure 2—video 1**. 13A neurons in the right front leg neuromere (T1R). Primary neurons in brown. Secondary neurons in green and red. Three sub-bundles of hemi-lineage 13A neurons are shown across 2D EM sections (left) and 3D rendering (right) of the cell bodies and 13A axonal tracks entering the neuropil.

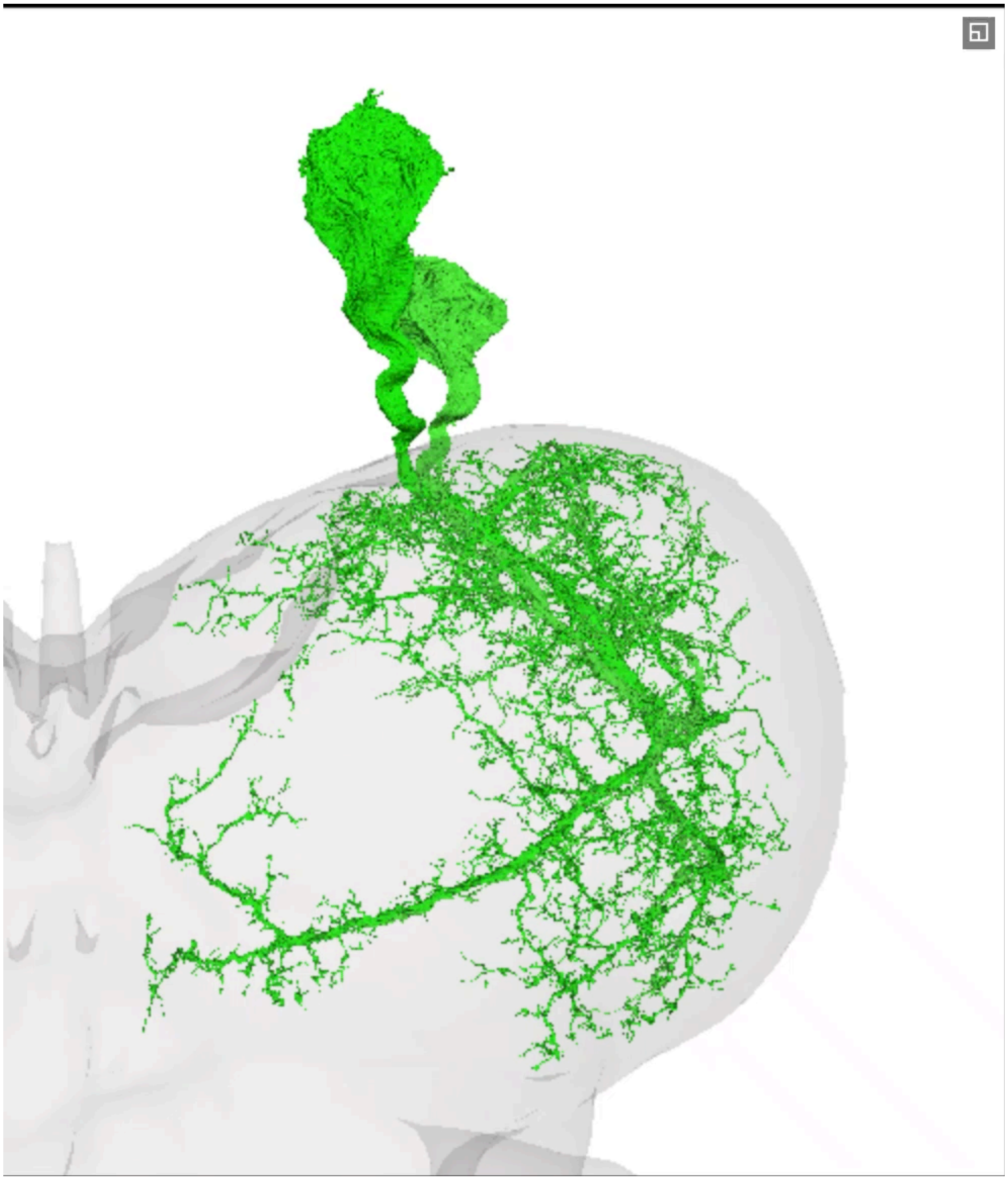

**Figure 2—video 2**. 13A morphological clusters

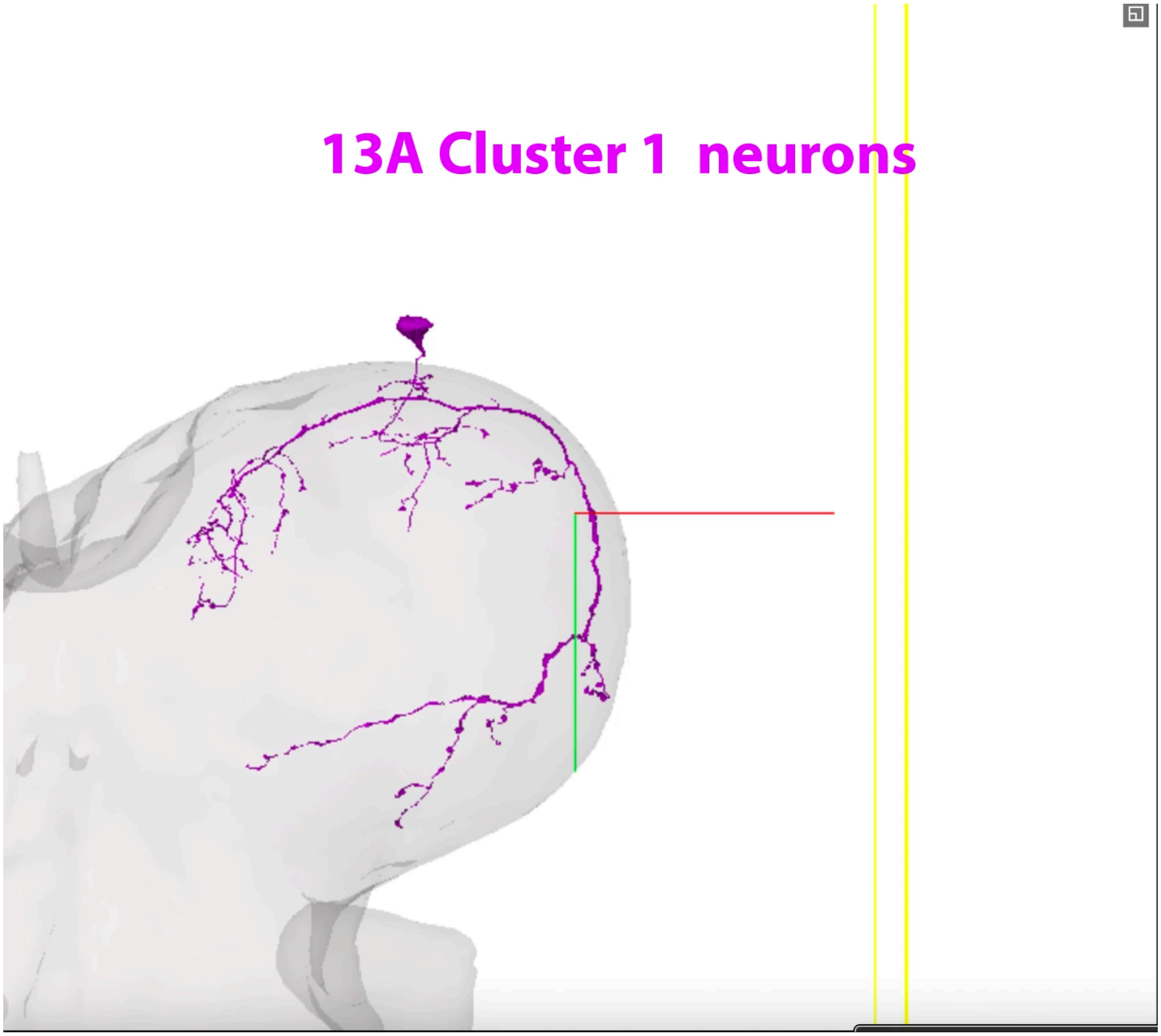

**Figure 2—video 3**. 13A cluster 6 neurons (red) and downstream Tibia extensor MNs (feti and seti, green)

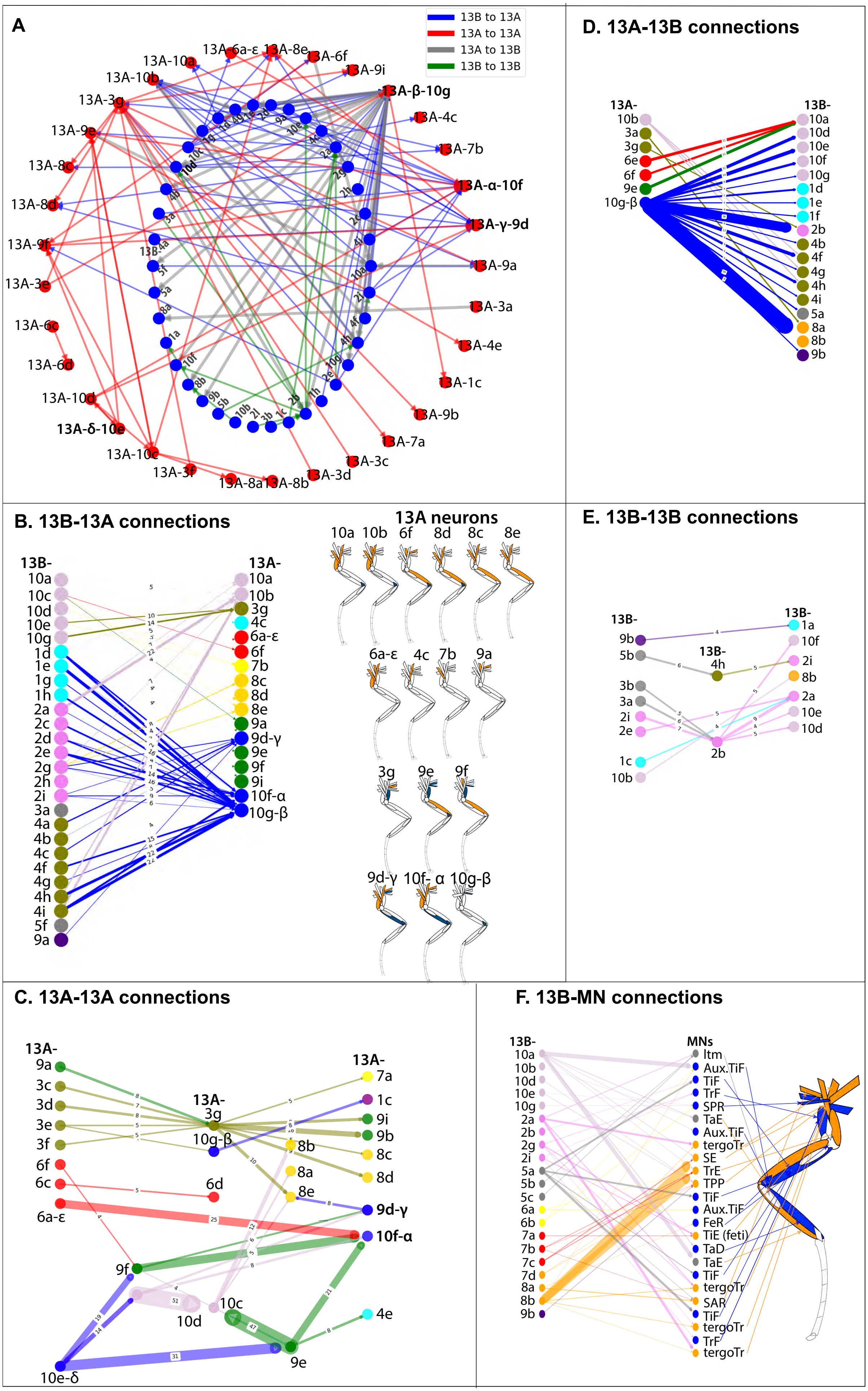
A
13B to 13A
13A to 13A
13A to 13B
13B to 13B
13A-10a
13A-10b
13A-3g
13A-9e
13A-8c
13A-8d
13A-9f
13A-3e
13A-6c
13A-6d
13A-10d
13A-δ-10e
13A-10c
13A-3f
13A-8a
13A-8b
13A-3d
13A-3c
13A-7a
13A-9b
13A-1c
13A-4e
13A-3a
13A-9a
13A-γ-9d
13A-α-10f
13A-7b
13A-4c
13A-β-10g
13A-9i
13A-6f
13A-6a-ε
13A-8e
D. 13A-13B connections
13A-
10b
3a
3g
6e
6f
9e
10g-β
13B-
10a
10d
10e
10f
10g
1d
1e
1f
2b
4b
4f
4g
4h
4i
5a
8a
8b
9b
B. 13B-13A connections
13B-
10a
10c
10d
10e
10g
1d
1e
1g
1h
2a
2c
2d
2e
2g
2h
2i
3a
4a
4b
4c
4f
4g
4h
4i
5f
9a
13A-
10a
10b
3g
4c
6a-ε
6f
7b
8c
8d
8e
9a
9d-γ
9e
9f
9i
10f-α
10g-β
13A neurons
10a 10b 6f 8d 8c 8e
6a-ε 4c 7b 9a
3g 9e 9f
9d-γ 10f- α 10g-β
E. 13B-13B connections
13B-
9b
5b
3b
3a
2i
2e
1c
10b
13B-
4h
2b
13B-
1a
10f
2i
8b
2a
10e
10d
C. 13A-13A connections
13A-
9a
3c
3d
3e
3f
6f
6c
6a-ε
13A-
3g
10g-β
6d
13A-
7a
1c
9i
9b
8c
8d
8b
8a
8e
9d-γ
10f-α
9f
10c
10d
10e-δ
9e
4e
F. 13B-MN connections
13B-
10a
10b
10d
10e
10g
2a
2b
2g
2i
5a
5b
5c
6a
6b
7a
7b
7c
7d
8a
8b
9b
MNs
ltm
Aux.TiF
TiF
TrF
SPR
TaE
Aux.TiF
tergoTr
SE
TrE
TPP
TiF
Aux.TiF
FeR
TiE (feti)
TaD
TaE
TiF
tergoTr
SAR
TiF
tergoTr
TrF
tergoTr

**Figure 3—figure supplement 1. Disinhibition Matrix**

**(A) Network Visualization of all 13A and 13B Neuronal Interconnections.** Network graph showing all 13B (presynaptic) to 13A (postsynaptic) connections (blue edges), 13A to 13A connections (red edges), 13A to 13B connections (gray edges) and 13B to 13B connections (green edges). 13A nodes in red. 13B nodes in blue.

**(B) Disinhibition mediated by 13B neurons**: A connectivity graph showing all 13B to 13A connections.The leg schematic on the right side shows targets of motor neurons inhibited by these 13A neurons, which are disinhibited by 13B neurons. Nodes of the same color within a lineage represent neurons within the same morphological cluster. Edge color corresponds to the postsynaptic 13A targets. Protractors/levators/extensors in orange and retractors, depressors, flexors in blue.

**(C) Disinhibition mediated by 13A-13A connections:** A connectivity graph highlighting connections within 13A neurons that leads to disinhibition of motor neurons. Nodes of the same color represent same 13A anatomical cluster. Primary 13A neuron nodes are highlighted in blue, except for 13AR-6a(-ε) that is highlighted in red. Color of the edges corresponding to the color of a presynaptic neuron.

**(D) Disinhibition mediated by 13A-13B connections:** A connectivity graph showing connections from 13A neurons to 13B neurons. Note that most of these postsynaptic 13B neurons are premotor (shown in panel F).

**(E) Disinhibition mediated by 13B-13B connections:** A connectivity graph showing interconnections between 13B neurons.

**(F) Premotor 13B neurons:** A connectivity graph showing premotor 13B neuron connections to MNs. 13B neurons that are morphologically related (same cluster) are shown in same color. 13B neurons that belong to the same cluster do not connect to same set of motor neurons. Protractors/levators/extensors in orange and retractors, depressors, flexors in blue.

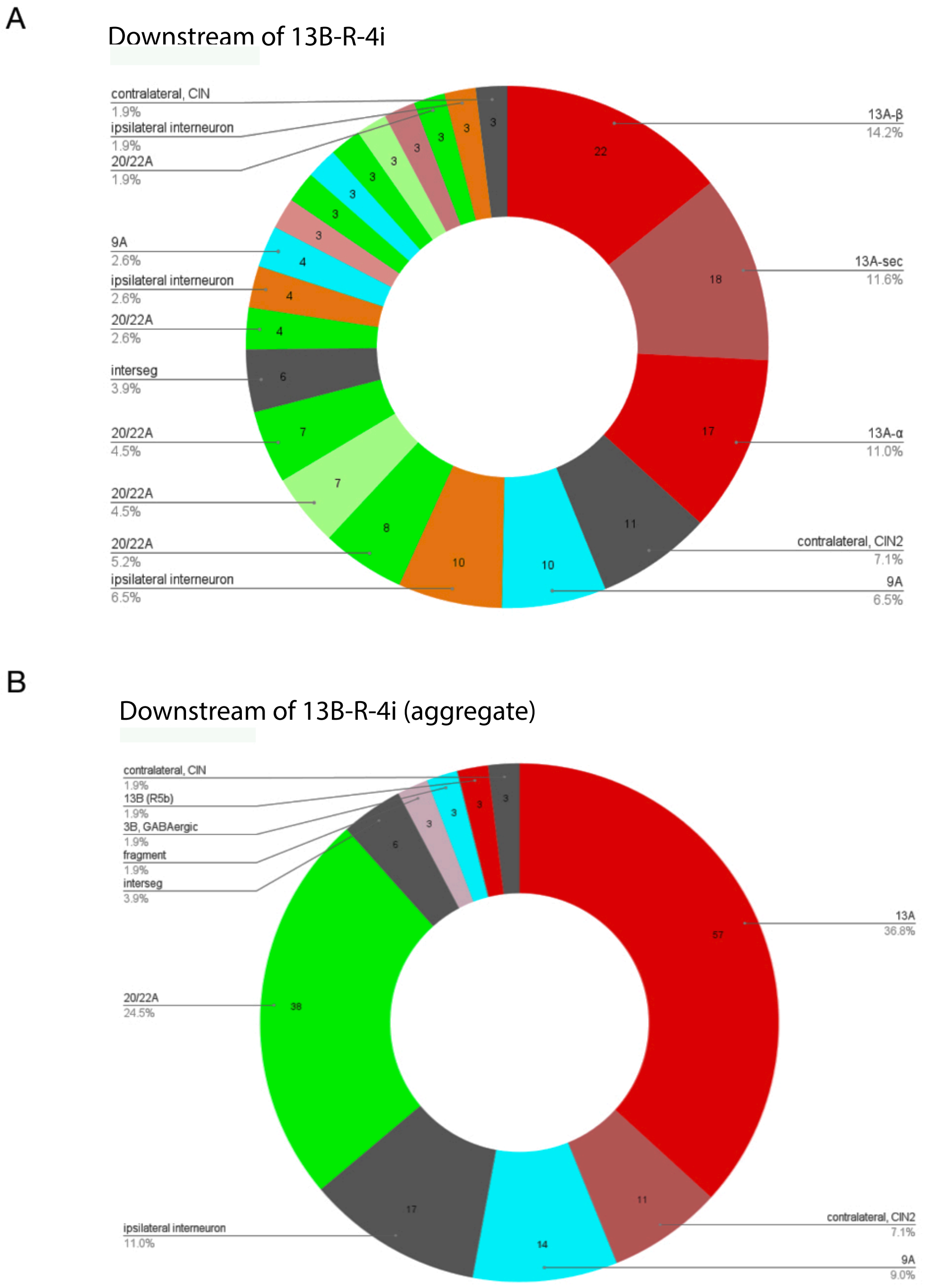

**Figure 3—figure supplement 2. Neurons Downstream of a 13B Neuron (13B-4i).** 13A neurons (red) comprise 36.8% of the total downstream synapses. 20/22A cholinergic neurons (green), 24.5% of the total synapses. 9A neurons (cyan)(6.5%), two contralateral interneurons (9%), 13B neuron (1.9%), 3B (1.9%), ipsilateral interneurons (possibly 21A?) (11%) of the total downstream partners. A. Individual downstream neurons. B. Aggregate of the same type of neurons.

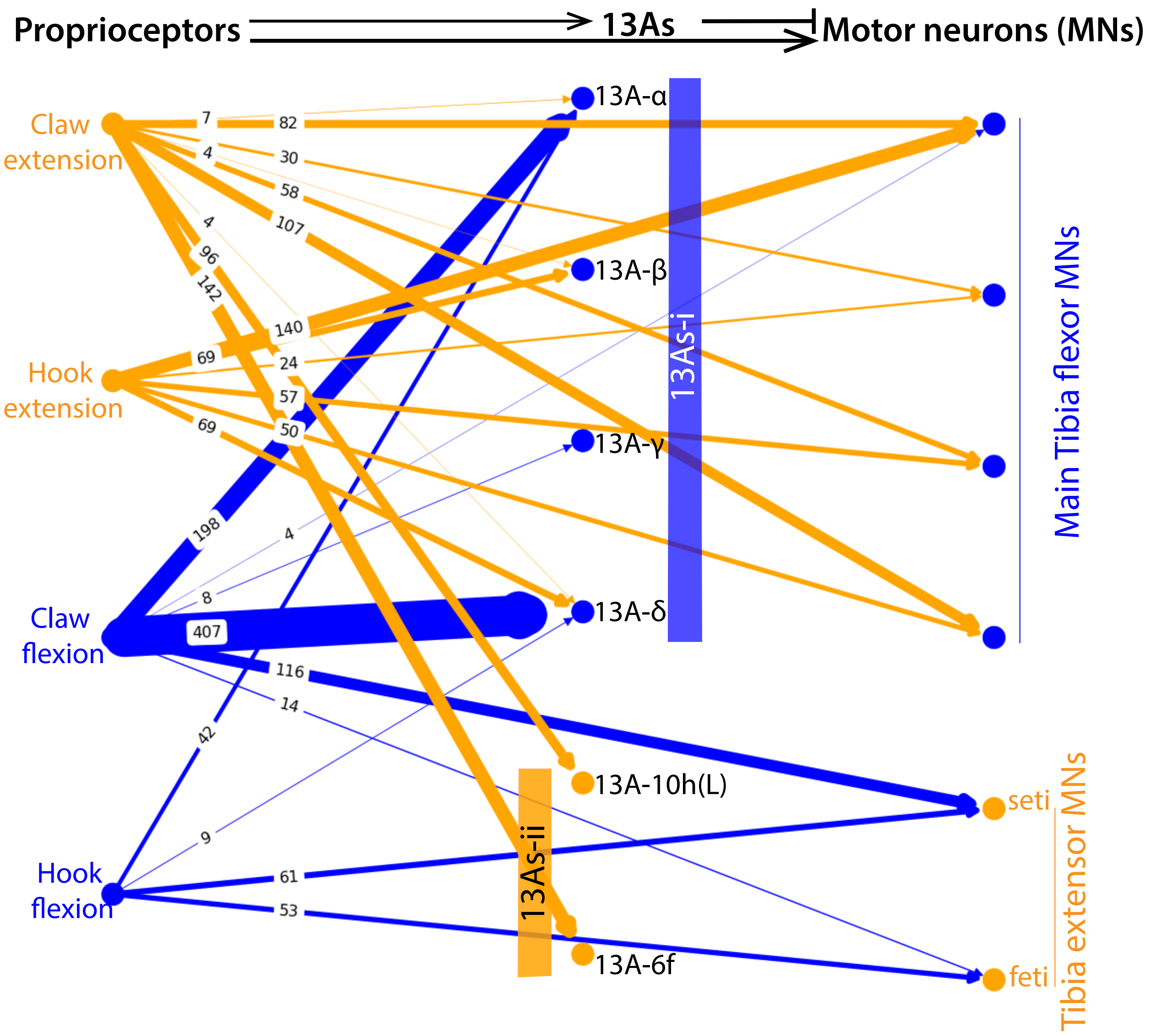

**Figure 3—Figure Supplement 3. Sensory Feedback Onto Inhibitory 13A Neurons and Motor Neurons.** Connectivity matrix showing position sensing claw neurons and motion sensing hook neurons send feedback connections to 13A neurons and antagonistic MNs. Flexion position and motion sensing proprioceptors (blue) neurons connect to tibia extensor MNs (orange) and primary 13A neurons (13As-i group) (blue). 13As-inhibit tibia flexor MNs. Thus, when the flexion is complete, it could induce extension and inhibit flexion via 13A neurons. Similarly, extension position and motion sensing proprioceptors (orange) neurons connect to tibia flexor MNs (blue) and two 13A neurons (13As-ii group) (orange) that inhibit tibia extensor MNs. Overall, extension position sensing neurons could activate flexion and inhibit extension. Synaptic weights are shown between connections also indicated by the edge thickness. Edge color corresponds to the identity of the proprioceptive neuron (Schematic in Figure 3D).

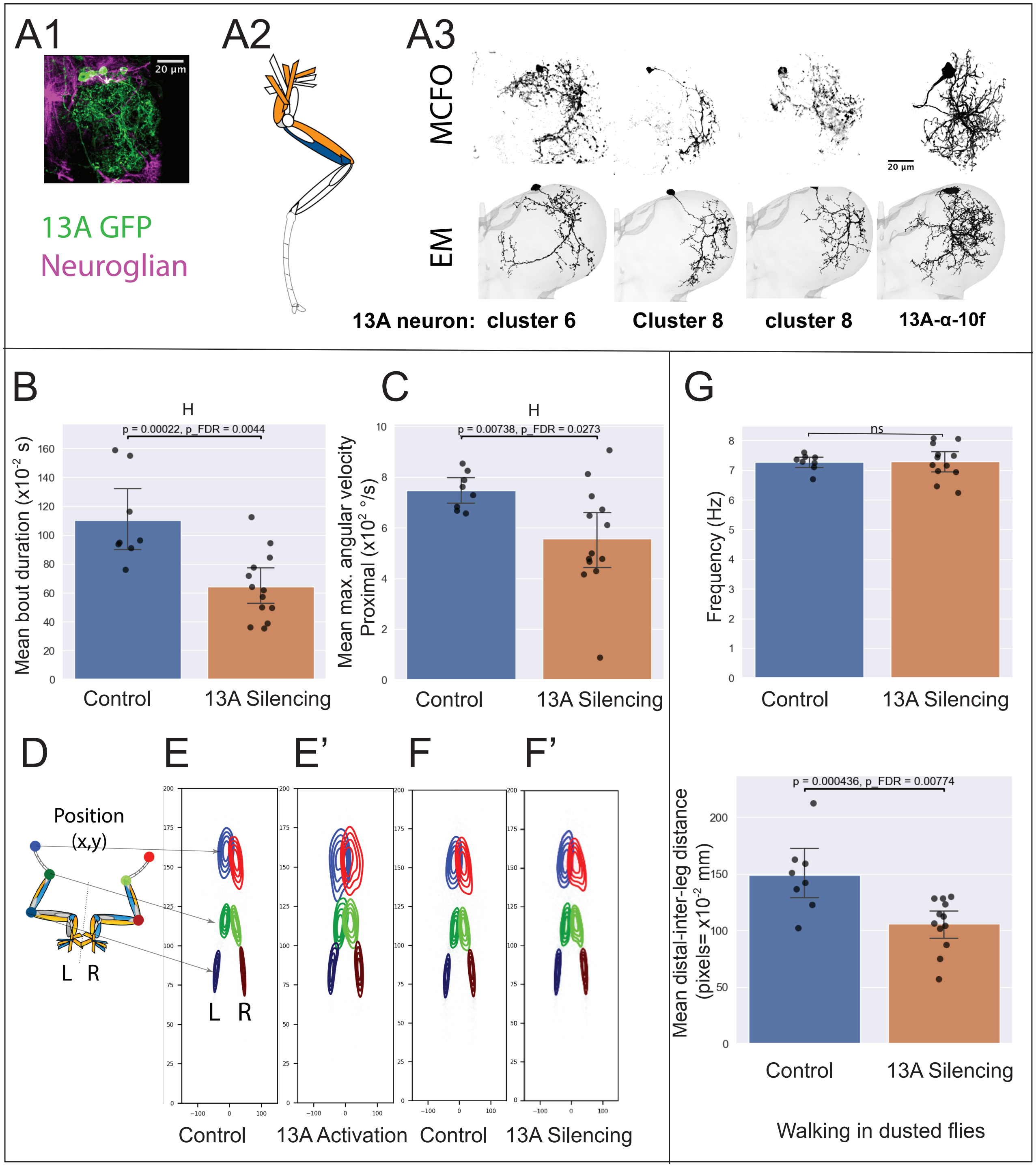

**Figure 4—figure supplement 1. 13A Neurons Regulate Leg Coordination During Grooming**

**Neuronal labeling and connectivity:** (A1) A confocal image showing six Dbx positive 13A neurons/hemisegment labeled by GFP driven by *R35G04-DBD, Dbx-AD Split GAL4.* T1 right front leg neuropil of the VNC is shown. GFP (green) labels 13A neurons. Neuroglian labels axon bundles in magenta. (A2) Schematic illustrating all muscles controlled by the motor neurons inhibited by specific 13A neurons. (A3) Top panel shows confocal images showing multicolor flip out clones (MCFO) (Nern et al., 2015) of 13A neurons. Bottom panel shows corresponding EM reconstructions of 13A neurons that resemble MCFO clones. We found four morphologically distinct targeted neurons among six 13A neurons. This may reflect incomplete labeling or morphological similarity between two neurons that can not be reliably distinguished. In the latter case, the two neurons likely belong to the same anatomical cluster hence connect to same set of MNs.

**Manipulating activity of Six Dbx positive 13A Neurons:** Silencing and activation of 13A neurons in dust-covered flies using *R35G04-DBD, Dbx-AD Split Gal4 > UAS Kir and UAS CsChrimson*, respectively. Control conditions include *AD-DBD Empty Split* for inactivation and *AD-DBD Empty Split with UAS CsChrimson* for activation. 13A inactivation (n=13 flies), activation (n=19 flies). Each panel compares control (blue) and experimental (orange) groups. Each dot represents the mean feature value for a single fly. Bars indicate the group mean, and whiskers represent the 95% confidence interval of the group mean. *P*-values (raw and FDR–corrected) are shown above each panel.

(B, C) Reduction in head grooming bout duration ($s*10^{-2}$) and maximum angular velocity ($° /s*10^{-2}$) of the proximal joint upon silencing of 13A neurons.

(D-F’) Joint Positions: Schematic showing position of various joints of the front legs is shown in F. Contour plots (probability distribution of joint positions) of the front legs of all the control and experimental flies during head grooming. The position of leg terminal (tarsus tip) is shown in colors: left leg in blue, right in red, the distal joint in dark and light green, and medial joint in dark blue and maroon. Joints positions are significantly altered upon activation of 13A neurons (E’), and slightly upon silencing (F’) of 13A neurons in dust-covered flies.

(G) Walking in dust-covered flies: Silencing 13A neurons did not affect the frequency of proximal joint movements (top panel), but reduced the distance between the tarsal tips of the front legs (bottom panel).

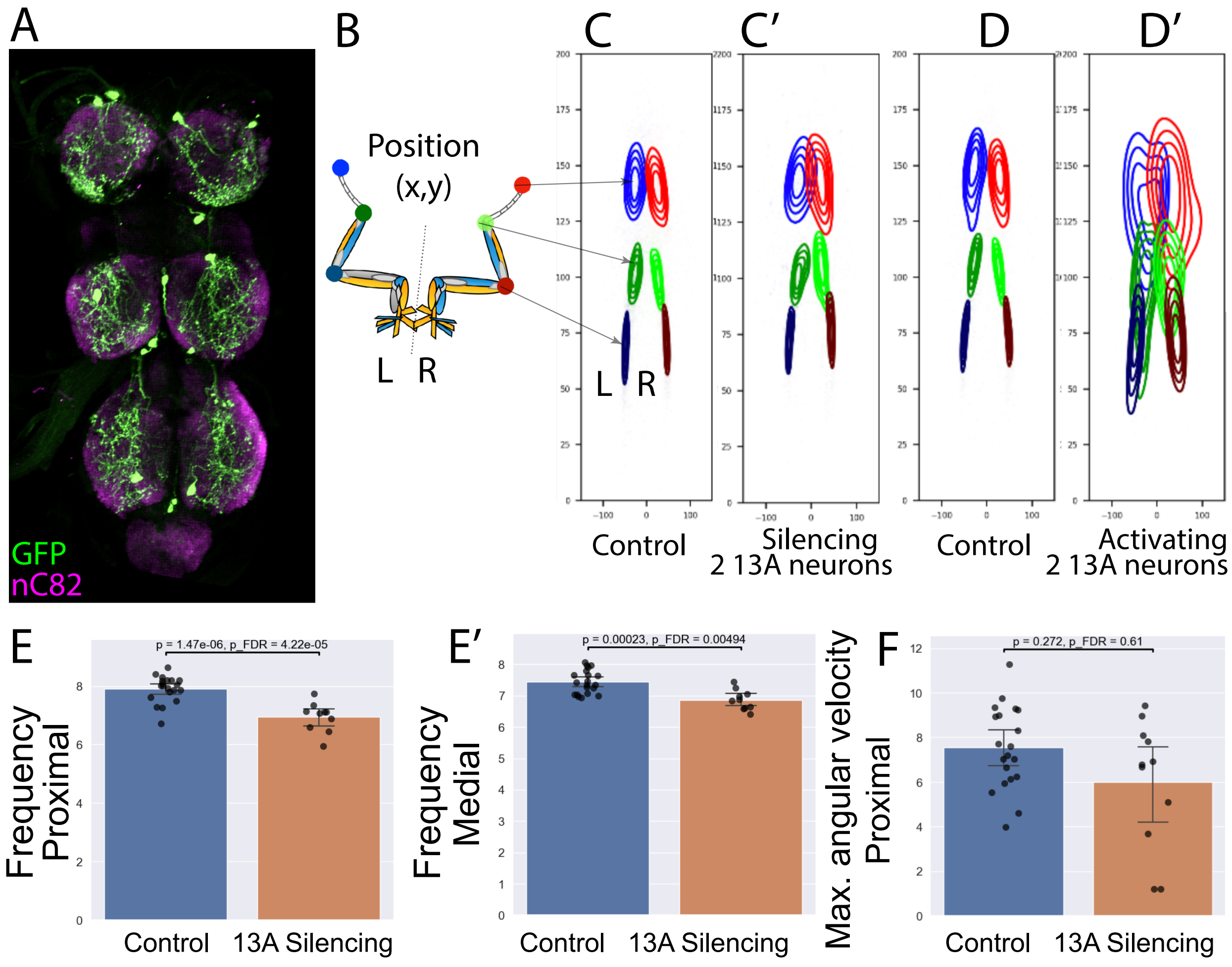

**Figure 4—figure supplement 2. Two Dbx Positive 13A Neurons Are Involved in Leg Coordination During Grooming in Dust-Covered Flies**

(A) Confocal image showing two Dbx positive 13A neurons/hemisegment labeled by GFP driven by *R11C07-DBD, Dbx-AD Split GAL4* in the adult VNC, labeled by GFP (green). nC82 (magenta) labels synaptic neuropil.

(B-G) Effects of manipulating activity of two 13A neurons: Silencing and activation experiments in dusted flies using *R11C07-DBD, Dbx-AD Split Gal4 > UAS GTACR1 and UAS CsChrimson*, respectively. Control conditions include *AD-DBD Empty Split* with *UAS GTACR1* for inactivation and with *UAS CsChrimson* for activation. 13A inactivation (n=11), activation (n=4). Bar plots in each panel compares control (blue) and experimental (orange) groups. Each dot represents the mean feature value for a single fly. Bars indicate the group mean, and whiskers represent the 95% confidence interval of the group mean. *P*-values (raw and FDR–corrected) are shown above each panel.

(C-D’) Contour plots (probability distribution of joint positions) of the front legs during grooming actions. Joints positions are significantly altered upon silencing (C’) and activation of 13A neurons (D’) in dust covered flies. Joint positions are shown during head sweeps (C-D’).

(E,E’) Median frequency (Hz) of the proximal and medial joints decreases upon silencing of two 13A neurons.

(F) Maximum angular velocity (° /s*$10^{-2}$) of the proximal joint does not significantly reduce upon silencing of two 13A neurons.

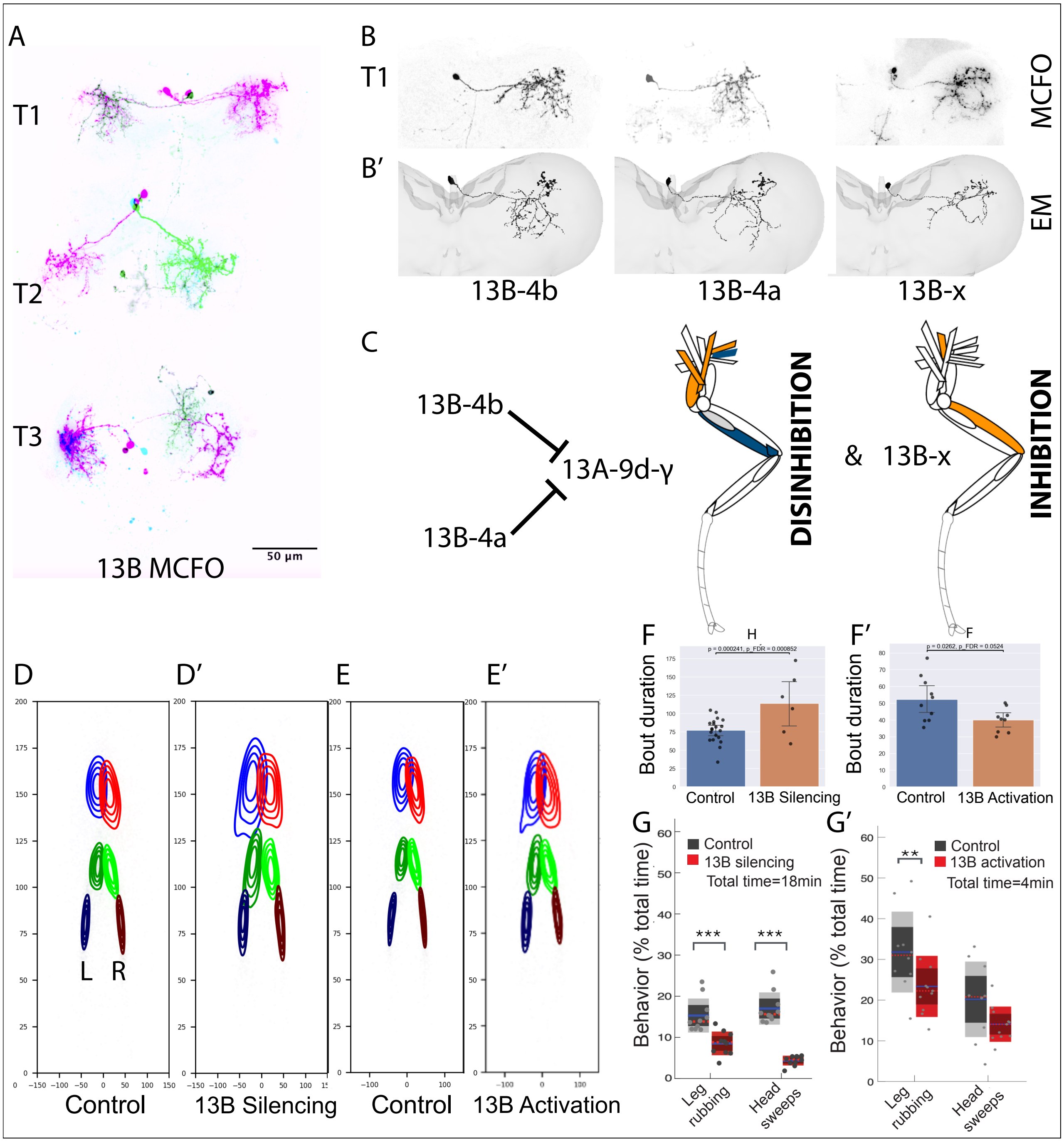

**Figure 4—figure supplement 3. Neuronal Labeling. Connectivity and Behavior of Specific 13B Neurons**

(A) Neuronal labeling: A confocal image showing clonal analysis of 2-3 13B neurons labeled in each hemisegment of the adult ventral nerve cord by *R11B07-DBD, GAD-AD Split GAL4.* Multicolor flip-out (MCFO) clones of all the 13B neurons labeled are shown in purple, green and black.

(B) MCFO clones of individual 13B neurons labeled in the right T1.

(B’) EM reconstructions of 13B neurons in right T1 that resemble these MCFO clones.

(C) Muscle targets disinhibited by two of these 13B neurons and those inhibited by one 13B neuron. Proximal (Th-C, C-Tr) extensor MNs and medial (Fe-Ti) flexor MNs are disinhibited while medial extensors are inhibited. Protractors/levators/extensors in orange and retractors, depressors, flexors in blue.

(D-E’) Contour plots (probability distribution of joint positions) of the front legs during grooming actions. Joints positions are significantly altered upon silencing (D’) and activation of 13B neurons (E’) in dust covered flies. Joint positions are shown during head sweeps (D-E’’).

(F, F’) Mean duration of head grooming bouts is longer upon silencing 13B neurons, whereas continuous activation of 13B neurons causes a slight, non-significant decrease in leg rubbing bouts in dusted flies. Bar plots compare control (blue) and experimental (orange) groups. Each dot represents the mean feature value for a single fly. Bars indicate the group mean, and whiskers represent the 95% confidence interval of the group mean. P-values (raw and FDR–corrected) are shown above each panel.

(G,G’) Percentage of time spent doing anterior grooming is reduced both upon continuous silencing and activation of 13B neurons. Box plots indicate the percentage of time dusted fly engaged in a given behavior. The solid blue line marks the mean, dark shading the 95% confidence interval, red dashed line the median, and light shading ±1 standard deviation. ***$P \leq 0.001$, **$P \leq 0.01$. Silencing experiment total time =18 min, scored with ABRS. Activation experiment total time =4 min, manually scored due to erratic leg extensions.

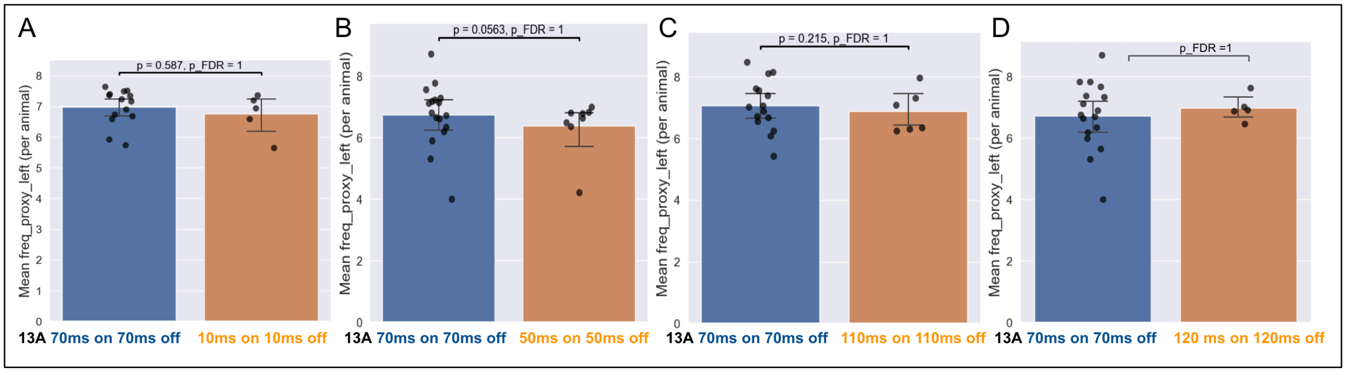

**Figure 5—figure supplement 1. Effect of varying optogenetic stimulation period on proximal joint cycle frequency.**

A) Comparison of mean proximal joint cycle frequency during 10 ms on/off stimulation (50 Hz) versus 70 ms on/off (~7 Hz).

(B) Comparison for 50 ms on/off (10 Hz) versus 70 ms on/off.

(C) Comparison for 110 ms on/off (~4.5 Hz) versus 70 ms on/off.

(D) Comparison for 120 ms on/off (~4 Hz) versus 70 ms on/off.

No significant differences were detected in any comparison (linear mixed model, p_FDR > 0.05). Bar plots in each panel compares control (~7Hz, blue) and experimental (orange) groups. Each dot represents the mean value for a single fly. Bars show group means, and whiskers represent the 95% confidence interval of the group mean. *P*-values (raw and FDR–corrected) are shown above each panel. These results indicate that pulsed activation triggers the circuit's intrinsic rhythm rather than pacing it at the stimulation frequency.

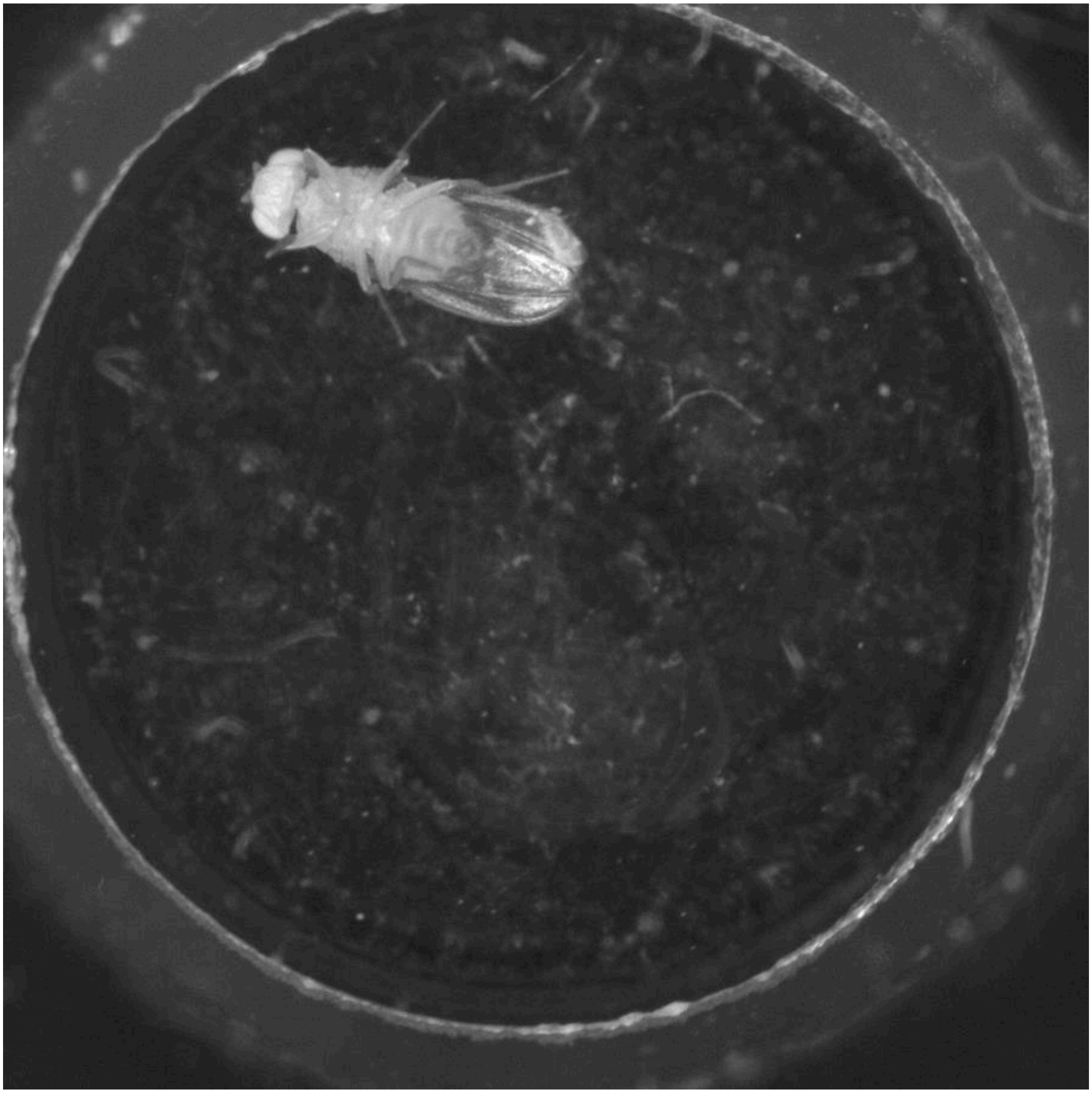

**<u>Figure 5—Video 1</u>** Activation of six 13A neurons with 70ms on and 70ms off pulses in undusted flies induces grooming and walking behaviors. In this video, light pulses start at t=30s with a pattern of 70ms on and 70ms off. Light is off from t=0-30s.

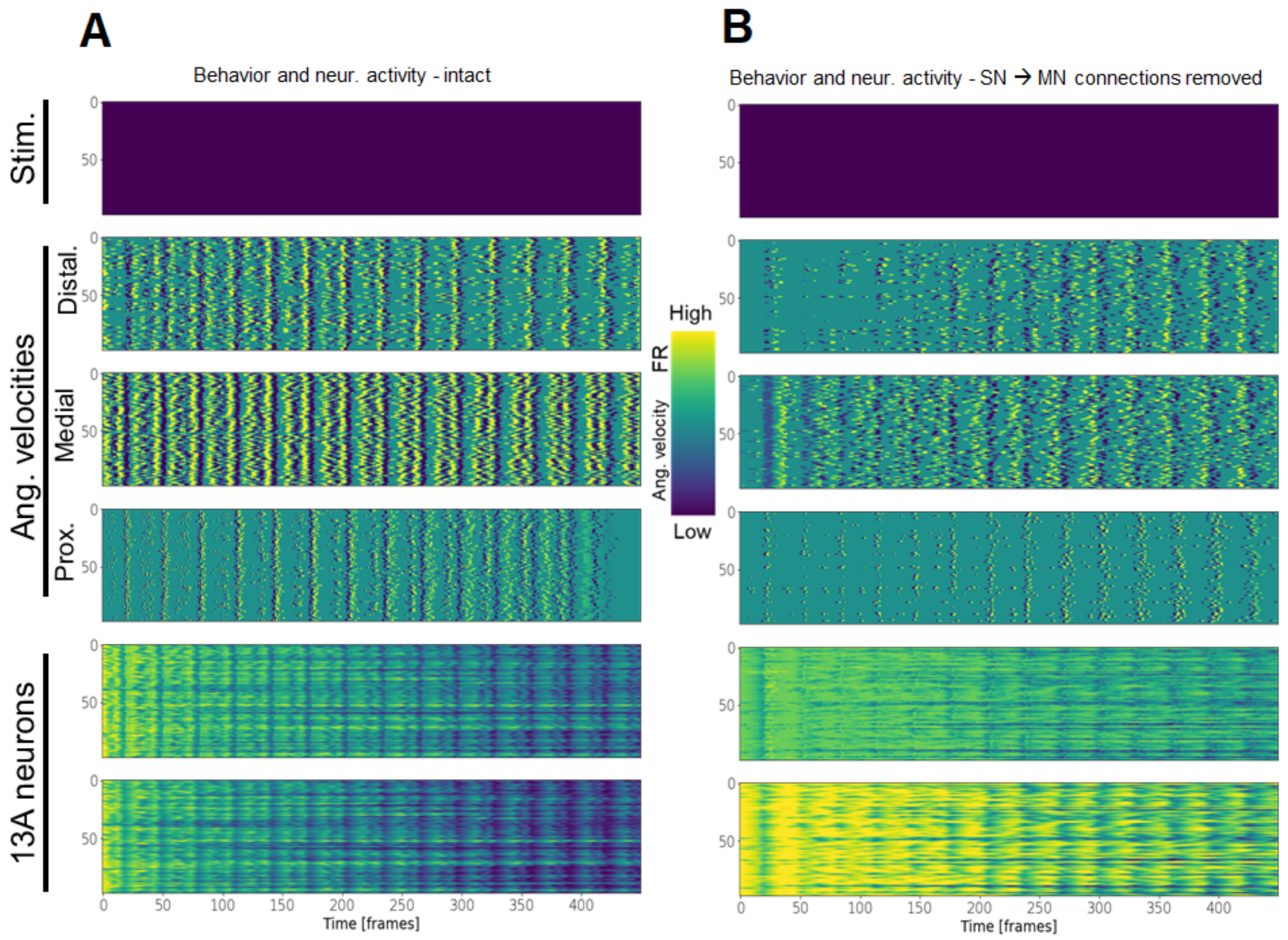

**Figure 6—Figure Supplement 1. Modeling the 13A Circuits.**

(A) Dynamics of angular velocities and 13A neurons, when no stimulus is given (Similar to Figure 7 J). (The simulation started running for 50 frames before Time=0 but it starts with very high peaks which were not plotted here for better visualization.)
(B) Same as A, but with proprioceptive SN → MN feedback connections obliterated.

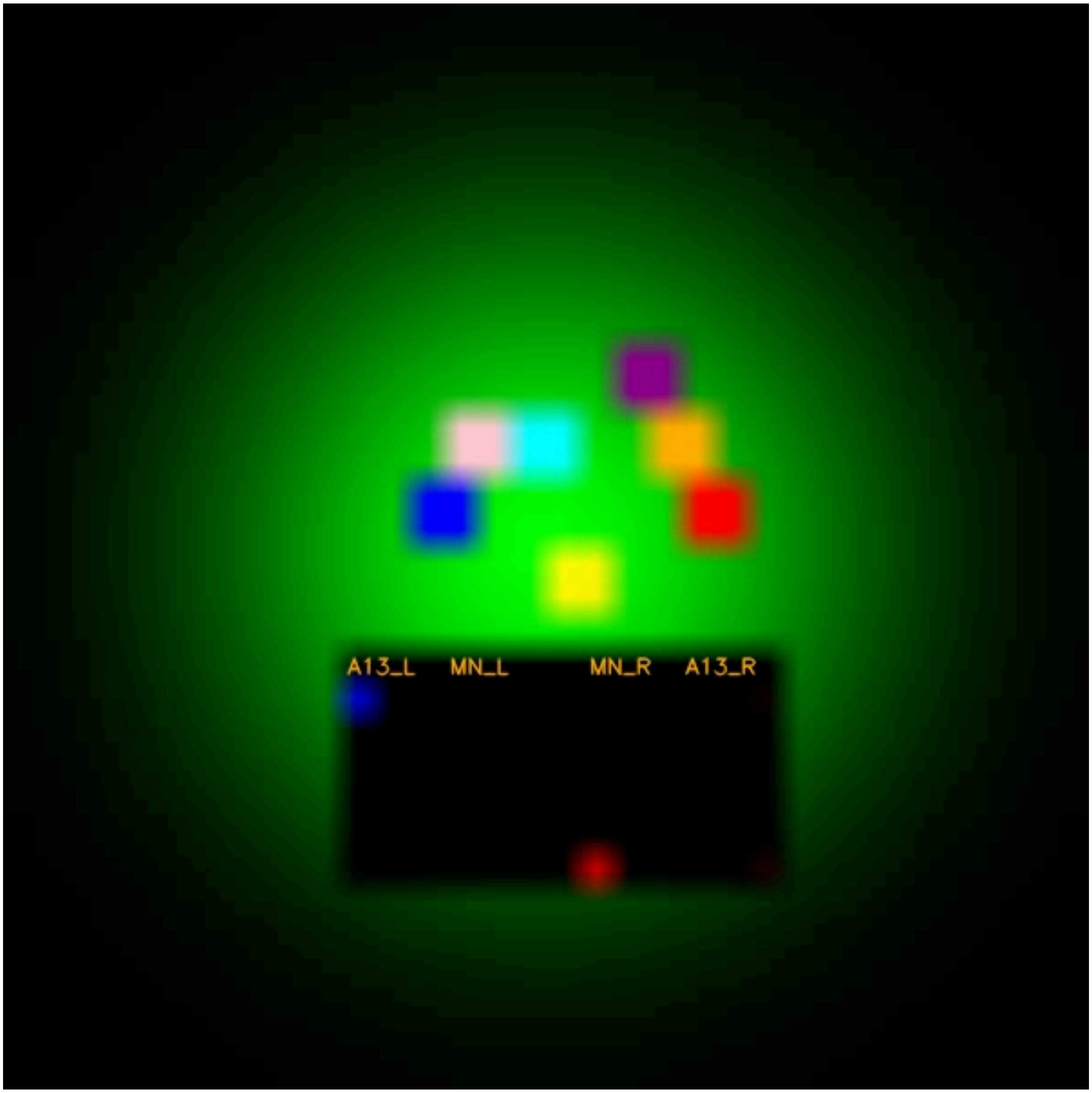

**Figure 6—Video 1.** **Modeling the 13A Circuits.** Description in Figure 6I.

| 13A neurons (Right T1) | FANC id | FANC coordinates | MANC type (Marin et al., Biorxiv) (2023) | MANC id |
|---|---|---|---|---|
| **13A-9d-γ** | 648518346483079845 | 51783, 91222, 3558 | IN13A001 | 10235 |
| **13A-10e-δ** | 648518346486953441 | 55683, 94740, 3977 | IN13A002 | 10820 |
| **13A-10f-α** | 648518346502652839 | 55113, 92627, 3760 | IN13A009 | 10230 |
| **13A-10g-β** | 648518346510045242 | 49361, 91971, 3631 | ~ IN13A003 | ~12376 |
| **13A-10a** | 648518346488772686 | 48080, 93048, 3688 | slightly ~IN13A012 | ~10553 |
| **13A-10b** | 648518346470640126 | 50638, 88360, 3240 | | |
| **13A-10c** | 648518346496914477 | 49195, 88089, 3275 | IN13A005 | 11770 |
| **13A-10d** | 648518346492366380 | 58416, 92888, 3866 | ~ IN13A015 | ~15715 |
| **13A-1a** | 648518346489892300 | 55164, 87638, 3027 | | |
| **13A-1b** | 648518346491127843 | 55377, 89872, 3431 | ~IN13A043 | ~30176 |
| **13A-1c** | 648518346490606940 | 50012, 90127, 3152 | | |
| **13A-1d** | 648518346504903795 | 58366, 89914, 3403 | | |
| **13A-1e** | 648518346490842877 | 55711, 89404, 3405 | | |
| **13A-1f** | 648518346511460661 | 52632, 89121, 3314 | | |
| **13A-2a** | 648518346489943859 | 54359, 90204, 3386 | IN13A071 | 172784 |
| **13A-2b** | 648518346481847711 | 53810, 91185, 3527 | | |
| **13A-3a** | 648518346496910338 | 48329, 88933, 3191 | | |
| **13A-3b** | 648518346507220040 | 48745, 86752, 3123 | | |
| **13A-3c** | 648518346495944043 | 53074, 88563, 3247 | | |
| **13A-3d** | 648518346493957489 | 59092, 91455, 3623 | | |
| **13A-3e** | 648518346459856132 | 57694, 92199, 3883 | | |
| **13A-3f** | 648518346488925169 | 53850, 88166, 3202 | | |
| **13A-3g** | 648518346499567451 | 54722, 90871, 3568 | | |
| **13A-3h** | 648518346501458787 | 47416, 87026, 3109 | | |
| **13A-3i** | 648518346502499667 | 53488, 87133, 3119 | ~ IN13A055 | ~31435 |
| **13A-3j** | 648518346481824415 | 51296, 87792, 3193 | ~IN13A047 | ~22393 |
| **13A-3k** | 648518346477176080 | 59399, 89215, 3309 | | |
| **13A-4a** | 648518346481645249 | 52182, 87862, 3083 | IN13A037 | 18827 |

| | | | | |
|---|---|---|---|---|
| **13A-4b** | 648518346502031978 | 55938, 89655, 3206 | | |
| **13A-4c** | 648518346475742644 | 50180, 87749, 3125 | | |
| **13A-4d** | 648518346490002483 | 50758, 91265, 3410 | | |
| **13A-4e** | 648518346499555931 | 48796, 89383, 3166 | | |
| **13A-5a** | 648518346509759363 | 58488, 92096, 3744 | IN13A060 | 23084 |
| **13A-5b** | 648518346475992546 | 50343, 87218, 3152 | | |
| **13A-5c** | 648518346515388743 | 55808, 91176, 3524 | IN13A041 | 21874 |
| **13A-5d** | 648518346488810830 | 57104, 89816, 3457 | IN13A060 | 16680 |
| **13A-5e** | 648518346488798392 | 57484, 91766, 3616 | | |
| **13A-5f** | 648518346486720303 | 50943, 90402, 3229 | | |
| **13A-5g** | 648518346496790140 | 54823, 88806, 3269 | | |
| **13A-5h** | 648518346490233883 | 51097, 89326, 3199 | | |
| **13A-6a-ε** | 648518346498563633 | 56616, 93624, 3918 | IN13A011 | 162586 |
| **13A-6b** | 648518346496931586 | 56616, 92824, 4050 | IN13A027 | 17619 |
| **13A-6c** | 648518346503236987 | 53992, 88752, 3163 | IN13A027 | 17252 |
| **13A-6d** | 648518346496223178 | 58635, 90741, 3546 | | ~19825 |
| **13A-6e** | 648518346491582906 | 59744, 88800, 3219 | ~IN13A042 | ~19825 |
| **13A-6f** | 648518346477206728 | 48760, 90632, 3495 | IN13A020 | 16744 |
| **13A-7a** | 648518346475595904 | 51067, 89384, 3298 | | |
| **13A-7b** | 648518346493912056 | 57282, 90706, 3505 | | |
| **13A-7c** | 648518346481935037 | 53489, 89742, 3404 | | |
| **13A-8a** | 648518346507214367 | 47821, 91396, 3352 | | |
| **13A-8b** | 648518346492957870 | 49102, 89285, 3418 | | |
| **13A-8c** | 648518346507315528 | 49397, 90429, 3418 | | |
| **13A-8d** | 648518346477160720 | 51624, 90320, 3439 | ~IN13A051 | ~101053 |
| **13A-8e** | 648518346494637032 | 49326, 89758, 3240 | | |
| **13A-9a** | 648518346521654521 | 47508, 88652, 3276 | IN13A050 | 31876 |
| **13A-9b** | 648518346496996404 | 49049, 92005, 3445 | | |
| **13A-9c** | 648518346518954719 | 54320, 87725, 3112 | ~IN13A035 | ~16080 |
| **13A-9e** | 648518346483398220 | 55912, 91368, 4070 | IN13A014 | 12790 |
| **13A-9f** | 648518346507449887 | 58219, 93783, 3911 | | |
| **13A-9g** | 648518346500177663 | 47886, 89723, 3334 | | |

| | | | |
|---|---|---|---|
| **13A-9h** | 648518346491573690 | 52349, 88228, 3203 | |
| **13A-9i** | 648518346524266501 | 50757, 88516, 3100 | |
| | | | ~ somewhat similar |
| **LINK R & L** | 13A right and Left | | |

| 13B (Right T1) | FANC ID | cell body coordinates |
|---|---|---|
| **13B-10a** | 648518346487050905 | 29608, 96504, 3357 |
| **13B-10b** | 648518346496337469 | 33624, 96848, 3441 |
| **13B-10c** | 648518346472698059 | 33032, 93768, 3319 |
| **13B-10d** | 648518346482001853 | 34904, 98024, 3432 |
| **13B-10e** | 648518346493383154 | 36624, 93656, 3314 |
| **13B-10f** | 648518346490249143 | 33056, 95648, 3415 |
| **13B-10g** | 648518346486447496 | 34000, 98264, 3320 |
| **13B-1a** | 648518346473301348 | 28462, 93468, 3173 |
| **13B-1b** | 648518346495825259 | 29612, 94321, 3266 |
| **13B-1c** | 648518346524174341 | 28242, 94354, 3181 |
| **13B-1d** | 648518346497088740 | 28092, 92452, 3172 |
| **13B-1e** | 648518346481492623 | 29186, 93423, 3245 |
| **13B-1f** | 648518346489719148 | 28457, 93932, 3304 |
| **13B-1g** | 648518346477208592 | 33378, 94661, 3356 |
| **13B-1h** | 648518346518812383 | 31525, 94153, 3238 |
| **13B-1i** | 648518346517337642 | 29000, 91552, 3131 |
| **13B-2a** | 648518346491055651 | 37806, 92227, 3272 |
| **13B-2b** | 648518346502643879 | 34607, 99358, 3478 |
| **13B-2c** | 648518346492013963 | 30112, 91667, 3096 |
| **13B-2d** | 648518346488885853 | 30810, 93887, 3273 |
| **13B-2e** | 648518346511706992 | 35618, 94709, 3364 |

| | | |
|---|---|---|
| **13B-2g** | 648518346491540392 | 36192, 93148, 3349 |
| **13B-2h** | 648518346507671417 | 30219, 92361, 3195 |
| **13B-2i** | 648518346531614746 | 34433, 93768, 3164 |
| **13B-2j** | 648518346486358930 | 35954, 95571, 3290 |
| **13B-2k** | 648518346486579360 | 30546, 90926, 3140 |
| **13B-2l** | 648518346502083434 | 32316, 92824, 3251 |
| **13B-2x** | 648518346499593307 | 35320, 93548, 3219 |
| **13B-3a** | 648518346511380098 | 34532, 97120, 3352 |
| **13B-3b** | 648518346504189792 | 31828, 93758, 3295 |
| **13B-3c** | 648518346478258157 | 29011, 92164, 3202 |
| **13B-4a** | 648518346502171977 | 36238, 95692, 3413 |
| **13B-4b** | 648518346474473581 | 34613, 95266, 3354 |
| **13B-4c** | 648518346498710256 | 35403, 95923, 3438 |
| **13B-4d** | 648518346484593155 | 32137, 94237, 3174 |
| **13B-4e** | 648518346510036794 | 33595, 95110, 3307 |
| **13B-4f** | 648518346496723324 | 29822, 93229, 3230 |
| **13B-4g** | 648518346510015034 | 33157, 93079, 3227 |
| **13B-4h** | 648518346486377058 | 30561, 93180, 3167 |
| **13B-4i** | 648518346494421874 | 29334, 90576, 3099 |
| **13B-4j** | 648518346488235550 | 31496, 92024, 3192 |
| **13B-5b** | 648518346487880002 | 34720, 98730, 3515 |
| **13B-5c** | 648518346489590725 | 30272, 95432, 3382 |
| **13B-5d** | 648518346489713772 | 31040, 96408, 3343 |
| **13B-5e** | 648518346476010466 | 28936, 94968, 3262 |
| **13B-5f** | 648518346479581403 | 32440, 92056, 3236 |
| **13B-5g** | 648518346496414269 | 34248, 93712, 3324 |
| **13B-5a** | 648518346488443631 | 36146, 94512, 3237 |
| **13B-6a** | 648518346483420350 | 35720, 98288, 3469 |

| | | |
|---|---|---|
| **13B-6b** | 648518346486357650 | 35476, 96698, 3464 |
| **13B-7a** | 648518346487047833 | 29792, 94176, 3141 |
| **13B-7b** | 648518346494016369 | 35224, 99176, 3410 |
| **13B-7c** | 648518346517501477 | 37864, 95976, 3308 |
| **13B-7d** | 648518346474731714 | 37488, 94352, 3360 |
| **13B-8a** | 648518346483038117 | 33872, 100552, 3614 |
| **13B-8b** | 648518346489995863 | 38720, 92192, 3220 |
| **13B-9a** | 648518346489035242 | 32656, 94800, 3287 |
| **13B-9b** | 648518346496206438 | 32328, 95784, 3321 |
| **13B-9c** | 648518346500201215 | 33200, 96408, 3311 |
| 13B-x | 648518346499846213 | 36640, 96472, 3387 |
| 13B-X4 | 648518346500534332 | 37775, 95047, 3482 |
| 13B-X6 | 648518346481737921 | [31528, 96264, 3440] |
| 13B-X7 | 648518346481077338 | 31336, 92760, 3266 |
| 13B-x8 | 648518346489982771 | 31230, 94887, 3313 |

| **13A (T1 Left)** | **ID** | **coordinates** |
|---|---|---|
| **13A-α** | 648518346500132614 | 24048, 93712, 3295 |
| **13A-β** | 648518346488838734 | 19600, 93632, 3325 |
| **13A-γ** | 648518346484661507 | 20360, 91608, 3178 |
| **13A-δ** | 648518346472722185 | 14392, 92936, 3157 |
| **13A-10b** | 648518346490196954 | 11993, 96131, 3206 |
| **13A-10c** | 648518346496087564 | 15440, 90440, 2896 |
| **13A-10d** | 648518346517571364 | 20640, 90256, 3025 |
| **13A-10e** | 648518346487847147 | 26480, 92792, 3168 |
| **13A-10f** | 648518346488990961 | 24120, 93088, 3163 |
| **13A-10g** | 648518346477777628 | 14312, 92192, 3010 |
| **13A-10h** | 648518346491793832 | 26288, 94336, 3338 |
| **13A-1a** | 648518346517685352 | 23448, 91080, 2938 |

| 13A (T1 Left) | ID | coordinates |
|---|---|---|
| 13A-1b | 648518346481678785 | 22904, 92040, 3154 |
| 13A-2a | 648518346497137068 | 25416, 92152, 3026 |
| 13A-2b | 648518346520616246 | 25320, 90816, 3143 |
| 13A-2c | 648518346495795819 | 25112, 91168, 3016 |
| 13A-2d | 648518346501310230 | 26800, 89112, 3011 |
| 13A-2e | 648518346496635650 | 27968, 90056, 3090 |
| 13A-2f | 648518346498525522 | 26168, 89584, 3047 |
| 13A-2g | 648518346480238139 | 27160, 94768, 3286 |
| 13A-3a | 648518346480464854 | 27280, 92072, 3106 |
| 13A-3b | 648518346486759395 | 25184, 92936, 3089 |
| 13A-3c | 648518346496505212 | 24136, 91152, 3031 |
| 13A-3d | 648518346489955788 | 22707, 92458, 2894 |
| 13A-3e | 648518346489955788 | 22584, 92112, 2938 |
| 13A-3g | 648518346488710840 | 15496, 95368, 3409 |
| 13A-4a | 648518346497975655 | 25344, 91616, 3099 |
| 13A-4b | 648518346498584881 | 19048, 91616, 3033 |
| 13A-4c | 648518346487654612 | 22128, 90520, 2913 |
| 13A-4d | 648518346503987240 | 18200, 91224, 2997 |
| 13A-4e | 648518346482173076 | 18200, 91280, 3108 |
| 13A-5a | 648518346499451995 | 17320, 91016, 3019 |
| 13A-5b | 648518346503984680 | 25656, 92024, 3195 |
| 13A-5c | 648518346504048078 | 22216, 91328, 2984 |
| 13A-5d | 648518346500110915 | 19272, 90032, 2944 |
| 13A-5e | 648518346497071532 | 24760, 90064, 3066 |
| 13A-6-ε | 648518346501484899 | 17272, 93616, 3314 |
| 13A-6a | 648518346486939545 | 22016, 89800, 2979 |
| 13A-6b | 648518346494486917 | 13072, 96624, 3346 |

| 13A (T1 Left) | ID | coordinates |
|---|---|---|
| 13A-6c | 648518346480509142 | 16288, 93240, 3200 |
| 13A-6d | 648518346496880180 | 19008, 90376, 3020 |
| 13A-6e | 648518346498076826 | 16056, 91864, 3106 |
| 13A-6f | 648518346491107107 | 13406, 95317, 3281 |
| 13A-7a | 648518346486288482 | 16160, 91072, 3025 |
| 13A-7b | 648518346497160600 | 17376, 92256, 3201 |
| 13A-7c | 648518346473216868 | 21136, 91840, 2978 |
| 13A-8a | 648518346490517084 | 23328, 91256, 3163 |
| 13A-8b | 648518346494588181 | 22264, 92496, 3250 |
| 13A-8c | 648518346479107791 | 15211, 91401, 2990 |
| 13A-9a | 648518346491531718 | 24144, 92464, 3027 |
| 13A-9b | 648518346499795084 | 25168, 90208, 2944 |
| 13A-9c | 648518346486941692 | 28176, 91208, 3159 |
| 13A-9d | 648518346502646183 | 26429, 92938, 3243 |
| 13A-9e | 648518346500029439 | 26720, 90856, 3149 |
| 13A-9f | 648518346493247730 | 25832, 90424, 3032 |
| 13A-9g | 648518346503131515 | 27056, 91952, 3212 |
| 13A | 648518346476966088 | 21005, 92746, 3160 |
| 13A | 648518346486963171 | 20384, 89590, 2927 |
| 13A | 648518346489565548 | 23950, 90091, 2913 |
| 13A | 648518346479948517 | 27753, 93259, 3244 |
| 13A | 648518346496562517 | 25303, 93564, 3164 |
| 13A | 648518346484458791 | 24178, 91813, 3112 |
| 13A | 648518346508904255 | 17317, 92205, 3085 |

**Table S1: 13A and 13B Neurons in the Front Leg Neuromere**
